# Participation of the rostral anterior cingulate cortex in defensive behavior induced by electrical stimulation of the dorsal periaqueductal gray and contextual fear conditioning

**DOI:** 10.1101/872648

**Authors:** Bruno de Oliveira Galvão, Alberto Filgueiras, Silvia Maisonnette, Thomas E. Krahe, J. Landeira-Fernandez

**Author notes:** Correspondence: J. Landeira-Fernandez, Núcleo de Neuropsicologia Clínica e Experimental, Laboratório de Neurociência Comportamental, Departamento de Psicologia, Pontifícia Universidade Católica do Rio de Janeiro, Rua Marquês de São Vicente, 225, Rio de Janeiro, RJ, 22453-900, Brazil.

## Abstract

The rostral anterior cingulate cortex (rACC) is a critical brain structure related to defensive behavior. However, still unclear is whether the rACC also plays a role in defensive behavior induced by electrical stimulation of the dorsal periaqueductal gray (dPAG). In the present study, rats were implanted with electrodes into the dPAG to determine freezing and escape response thresholds after sham or bilateral electrolytic lesions of the rACC. The duration of freezing behavior that outlasted electrical stimulation of the dPAG was also measured. The next day, these animals were subjected to contextual fear conditioning using footshock as an unconditioned stimulus. Lesions of the rACC did not change aversive freezing and escape response thresholds but disrupted post-dPAG stimulation freezing. The lesions also disrupted defensive freezing behavior and analgesia in the formalin test in response to contextual cues previously associated with footshock. These results indicate that the rACC is involved in some but not all aspects of defensive behavior generated at the level of the dPAG. The rACC also appears to play an important role in contextual fear conditioning.

## 1. Introduction

Anxiety disorders represent a heterogeneous group of psychopathologies with different etiologies. They likely reflect dysfunction of the neural circuits that are responsible for organizing defensive behavior systems to deal with threatening stimuli in the external environment. Several animal models of anxiety have been developed to improve our understanding of these disorders.

A gradual increase in electrical stimulation of the dorsal portion of the periaqueductal gray (dPAG) in rats causes an initial defensive freezing posture accompanied by piloerection and exophthalmus at lower intensities. As stimulation continues, vigorous escape responses, such as jumping and running, appear at higher intensities [1]. After the termination of electrical stimulation of the dPAG at the escape threshold, the animals engage in a long-lasting freezing response, a phenomenon that has been termed post-dPAG stimulation freezing [2].

Contextual fear conditioning represents another animal model of anxiety. A rat is exposed to a distinctive environment where it was previously exposed to an aversive unconditioned stimulus, such as an electric footshock. In such circumstances, the most prominent behavioral outcome is freezing behavior [3, 4]. Contextual fear conditioning can also activate endogenous pain-control systems, such as conditioned analgesia [5]. Conditioned analgesia has been suggested to allow a threatened animal to engage in necessary defensive reactions without being disturbed by competing reactions triggered by noxious stimulus [6].

Defensive behavior induced by contextual fear conditioning and electrical stimulation of the dPAG appears to employ distinct neural circuitries. For example, electrolytic lesions of the dPAG do not disrupt contextual fear conditioning [7]. Conversely, lesions of the amygdaloid complex or ventral portion of the PAG (vPAG) jeopardize freezing in response to contextual cues previously associated with footshock but do not change the aversive thresholds determined by electrical stimulation of the dPAG [8, 9]. Moreover, muscimol-induced inactivation of the amygdaloid complex reduced conditioned freezing in response to contextual cues previously associated with footshock and post-dPAG stimulation freezing but did not affect freezing or escape responses triggered by electrical stimulation of the dPAG [10]. Finally, serotonergic systems appear to differentially regulate contextual and post-dPAG stimulation freezing [11].

The rostral anterior cingulate cortex (rACC) is a critical brain structure related to defensive behavior. This cortical structure is part of the neural circuitry responsible for regulating the processing of nociceptive stimuli. Considerable evidence indicates that the rACC is involved in the memory formation of aversive tasks [12, 13]. For example, Einarsson and Nader [14] reported that microinfusion of the *N*-methyl-D-aspartate (NMDA) receptor NR2B subunit antagonist Ro25-6981 in the rACC reduced the freezing response to contextual cues previously associated with footshock. Moreover, the rACC appears to participate in inhibitory mechanisms of nociceptive processing. Accordingly, electrical stimulation of the rACC produced an analgesia effect in the hot-plate and tail-flick tests [15]. Indeed, human neuroimaging data indicate that the rACC is closely associated with placebo analgesia [16].

However, still unknown is whether the rACC also participates in conditioned analgesia induced by contextual fear conditioning. Furthermore, the role of the rACC in defensive responses triggered by electrical stimulation of the dPAG is also unclear. Therefore, the purpose of the present study was to investigate these issues. Electrical thresholds for defensive freezing and escape responses triggered by dPAG stimulation were measured in rats that received sham or bilateral rACC electrolytic lesions. Freezing behavior was measured after the termination of electrical stimulation of the dPAG at the escape threshold. Sham and lesioned animals were also subjected to the contextual fear conditioning protocol to investigate whether the electrolytic lesions of the rACC reduce freezing and conditioned analgesia responses induced by contextual cues previously associated with footshock.

## 2. Materials and Methods

### 2.1. Animals

Male albino rats from the animal colony of the Psychology Department, Pontifícia Universidade Católica do Rio de Janeiro, were used as subjects. Room temperature was controlled (24 ± 1°C), and the light/dark cycle was maintained on a 12 h on/off cycle. The animals weighed 250-300 g at the beginning of the experiment. They were housed individually in Plexiglas cages and given free access to food and water throughout the experiment. All of the experimental protocols used in this study were in accordance with the university research ethics committee and Brazilian Society of Neuroscience and Behaviour Guidelines for the Care and Use of Laboratory Animals (SBNeC), which are based on the United States National Institutes of Health Guide for Care and Use of Laboratory Animals (revised 1996).

### 2.2. Surgery

All of the animals were implanted with a unilateral guide cannula made of stainless steel aimed at the dPAG. Under tribromoethanol anesthesia (250 mg/kg, i.p.), each animal was fixed in a Kopf stereotaxic frame and locally injected with lidocaine (20 mg/ml). The upper incisor bar was set 3.3 mm below the interaural line such that the skull was horizontal between bregma and lambda. The following coordinates were used for the implantation of the guide cannula aimed at the dPAG and electrolytic lesions of the rACC, according to the Paxinos and Watson [17] rat brain atlas: dPAG (anterior/posterior, +2.3 mm; medial/lateral, −1.7 mm; dorsal/ventral, −4.5 mm), rACC (anterior/posterior, +2.7 mm; medial/lateral, ± 0.5 mm; dorsal/ventral, +2.2 mm). The guide cannula was attached to the skull with acrylic resin and three stainless steel screws. A stylet that was the same length as the guide cannula was introduced inside the guide cannula to prevent obstruction. Animals that were assigned to the rACC lesion group received bilateral electrolytic lesions aimed at the rACC by passing an anodal current (1.0 mA, 10 s) through the electrodes (Plastic One, Roanoke, VA, USA). Animals that were assigned to the sham lesion group underwent identical procedures with the exception that no electrical current was delivered.

### 2.3. Apparatus

Electrical stimulation of the dPAG and contextual fear conditioning occurred in the same observational chamber (25 × 20 × 20 cm). The chamber was placed inside a sound-attenuating box. A red light bulb (25 W) was placed inside the box, and a video camera was mounted on the back of the observation chamber so that the animal’s behavior could be observed on a monitor placed outside the experimental chamber. A ventilation fan attached to the box supplied background noise of 78 dB (A scale). The floor of the observational chamber was composed of 15 stainless steel, 4 mm diameter rods spaced 1.5 cm apart (center-to-center) that were wired to a shock generator and scrambler (AVS, SCR04; São Paulo, Brazil). An interface with eight channels (Insight Instruments, Ribeirão Preto, Brazil) connected the shock generator to a computer, which allowed the experimenter to apply an electric footshock. Ammonium hydroxide solution (5%) was used to clean the chamber before and after each subject.

### 2.4. Procedure

#### 2.4.1. Defensive behavior induced by dPAG electrical stimulation

One week after the surgery, each animal was placed inside the observational chamber. Five minutes later, aversive freezing and escape thresholds were determined using electrical stimuli (alternating current, 60 Hz, 20 s) presented through an electrical stimulator (AC, 60Hz, 15s) presented through a removable electrode (Plastics One) connected to a guide cannula aimed at the dPAG. The electrical stimulation was presented at 1 min intervals, with the current intensity increasing in 5 μA steps for measurements of aversive thresholds. The freezing threshold was operationally defined as the lowest current intensity that produced immobility, which was defined as the total absence of movement of the body or vibrissa, with the exception of movement required for respiration. The lowest current intensity that produced running (i.e., galloping movements) or jumping was considered the escape threshold. Animals with an escape threshold above 200 μA were discarded from the study. After reaching the escape threshold, the electrical stimulation of the dPAG stopped, and the animal remained in the observational chamber for an additional 12 min without any stimulation. During this period, freezing was scored using a time-sample procedure. Every 2 s, the animal’s freezing behavior was scored by a well-trained observer.

#### 2.4.2. Contextual fear conditioning and formalin test

One day after the end of electrical stimulation of the dPAG, all of the animals were subjected to the contextual fear conditioning paradigm and formalin test. The protocol consisted of training and test sessions. During the training session, each animal was placed in the observational chamber for 5 min. At the end of this period, three unsignaled 0.6 mA electric footshocks were delivered, with each shock lasting 1 s and an intershock interval of 20 s. The animal was returned to its home cage 3 min after the last shock. The test session occurred approximately 24 h after the training session. Before being placed in the observational chamber where the three shocks were delivered on the previous day, each rat was given a 0.05 ml subcutaneous injection of 15% formalin. The same time-sample procedure described above was used to score freezing and formalin-induced behavior (i.e., any licking or contact of the injected paw with the animal’s mouth or lifting and maintaining the injected paw off the grid floor for 60 min.

### 2.5. Histology

At the end of the experiment, the animals were deeply anaesthetized with chloral hydrate and intracardially perfused with a 0.9% saline solution followed by a 10% formalin solution. The cannula was removed, and the brain was placed in a 10% formalin solution. Three days later, the brain was frozen, and 50 μm brain sections were cut using a cryostat and stained with Cresyl blue to localize the cannula placements and lesion locations.

## 3. Results

All of the animals included in the analysis of the present study met the criteria for electrode placement in the dPAG and bilateral electrolytic lesions of the rACC. Histological examination of the brain slices indicated that all of the electrode tips were located inside the dPAG. Electrolytic lesions of the rACC were bilaterally symmetrical. Lesions included a cavity in the center of the lesion plus a region of chromatolysis that surrounded the cavity. Fig. 1 depicts the largest and smallest rACC lesions and a representative histological section of the electrolytic lesion. The final group samples were the following: rACC lesion (*n* = 10), sham lesion (*n* = 10).

**Figure 1.**
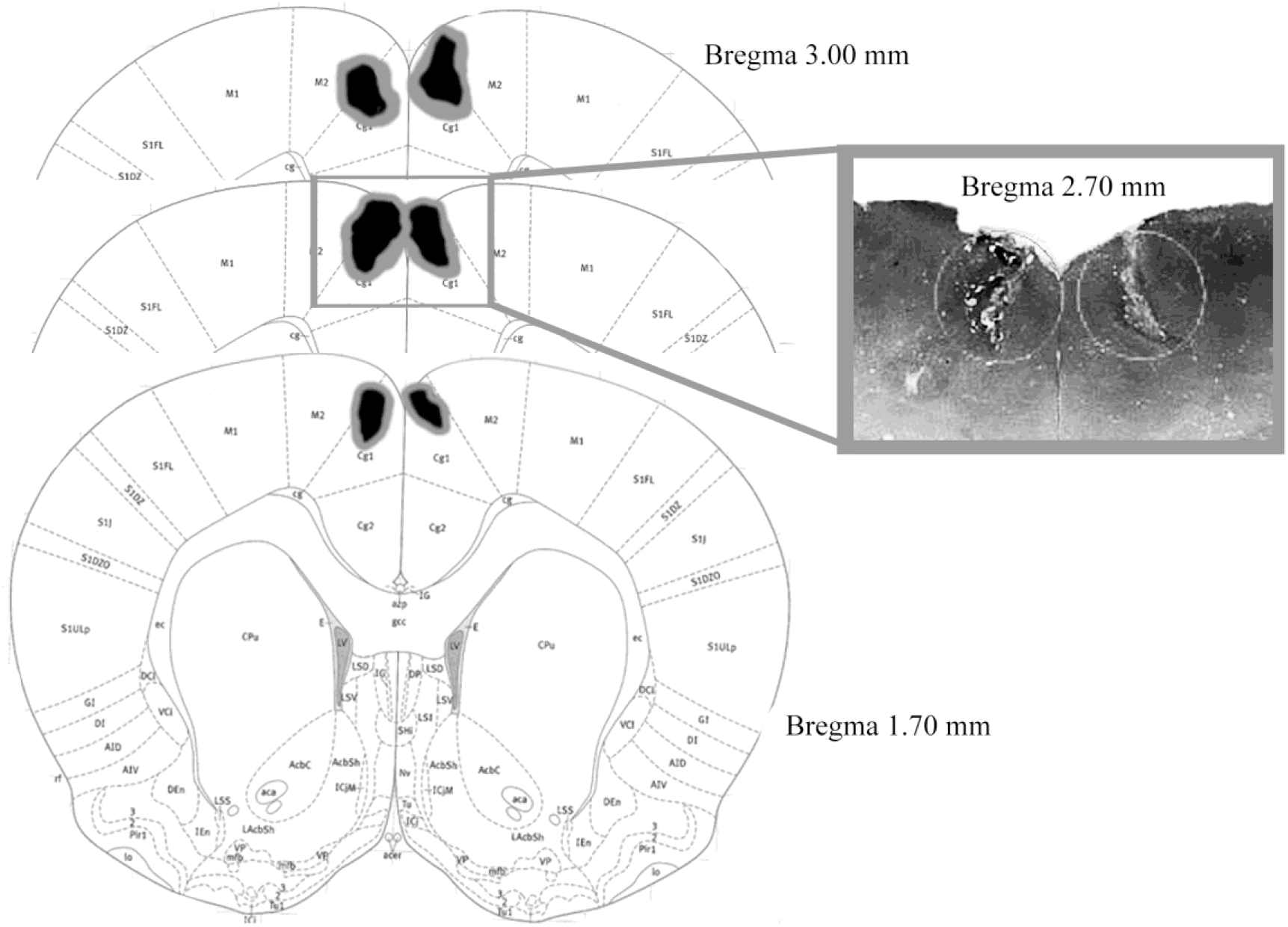
Representations of the largest and smallest rACC lesions depicted on images that represent slices at bregma +1.70, +2.70, and +3.00 mm. The photomicrograph depicts a representative coronal section through the rACC at bregma +2.70 mm.

As we reported previously [18], freezing and escape responses induced by electrical stimulation of the dPAG occurred in a stepwise fashion. As the intensity of the current applied to the dPAG increased, the animals suddenly stopped and became immobile, accompanied by piloerection and exophthalmus. At higher current intensities, this freezing behavior was followed by vigorous running and jumping reactions. The escape response stopped as soon as electrical stimulation of the dPAG was stopped. Fig. 2 depicts the mean and standard error of the mean (SEM) of the freezing and escape thresholds between the rACC and sham lesion groups. A two-way repeated-measures analysis of variance (ANOVA) was used to evaluate differences in aversive thresholds. The treatment (rAAC and sham lesion) was considered the between-subjects factor, and aversive threshold (freezing and escape) was considered the within-subjects factor. The 2 × 2 repeated-measures ANOVA revealed no treatment × aversive threshold interaction (*F*_1,18_ = 0.02; *p* = 0.91). No main effect of treatment was found (*F*_1,18_ = 0.01; *p* = 0.92). However, a significant main effect of aversive threshold was found (*F*_1,18_ = 42.27; *p* < 0.001).

**Figure 2.**
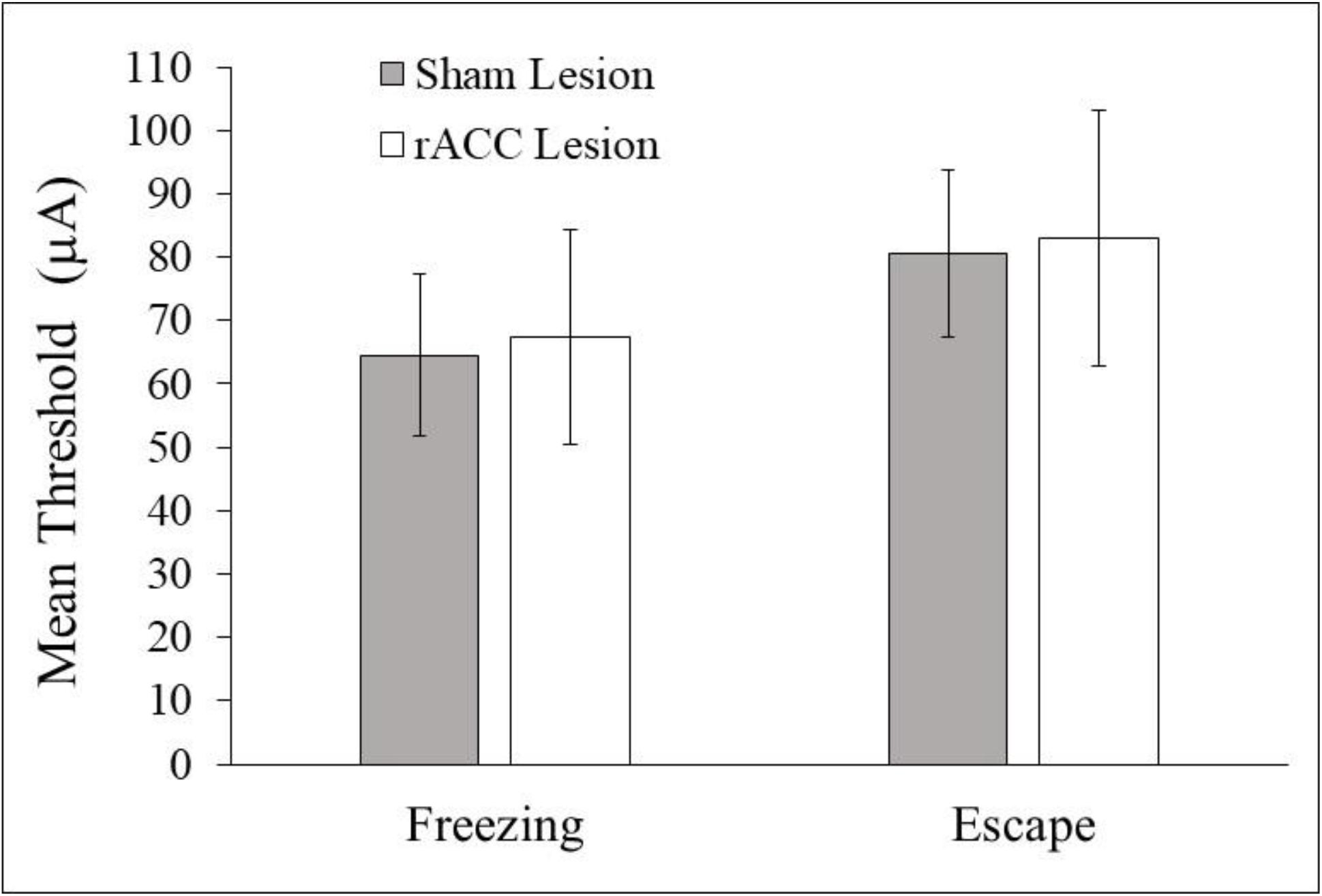
Mean (± SEM) freezing and escape thresholds induced by electrical stimulation of the dPAG in sham- and rACC-lesioned animals.

Fig. 3 shows the mean (± SEM) percentage of time that sham- and rACC-lesioned animals spent freezing after stimulation of the dPAG at the escape threshold. Student’s *t*-test indicated that rACC-lesioned animals exhibited less post-dPAG stimulation freezing behavior compared with sham-lesioned animals during the 12-min test period (*t*_18_ = 5.8; *p* < 0.001).

**Figure 3.**
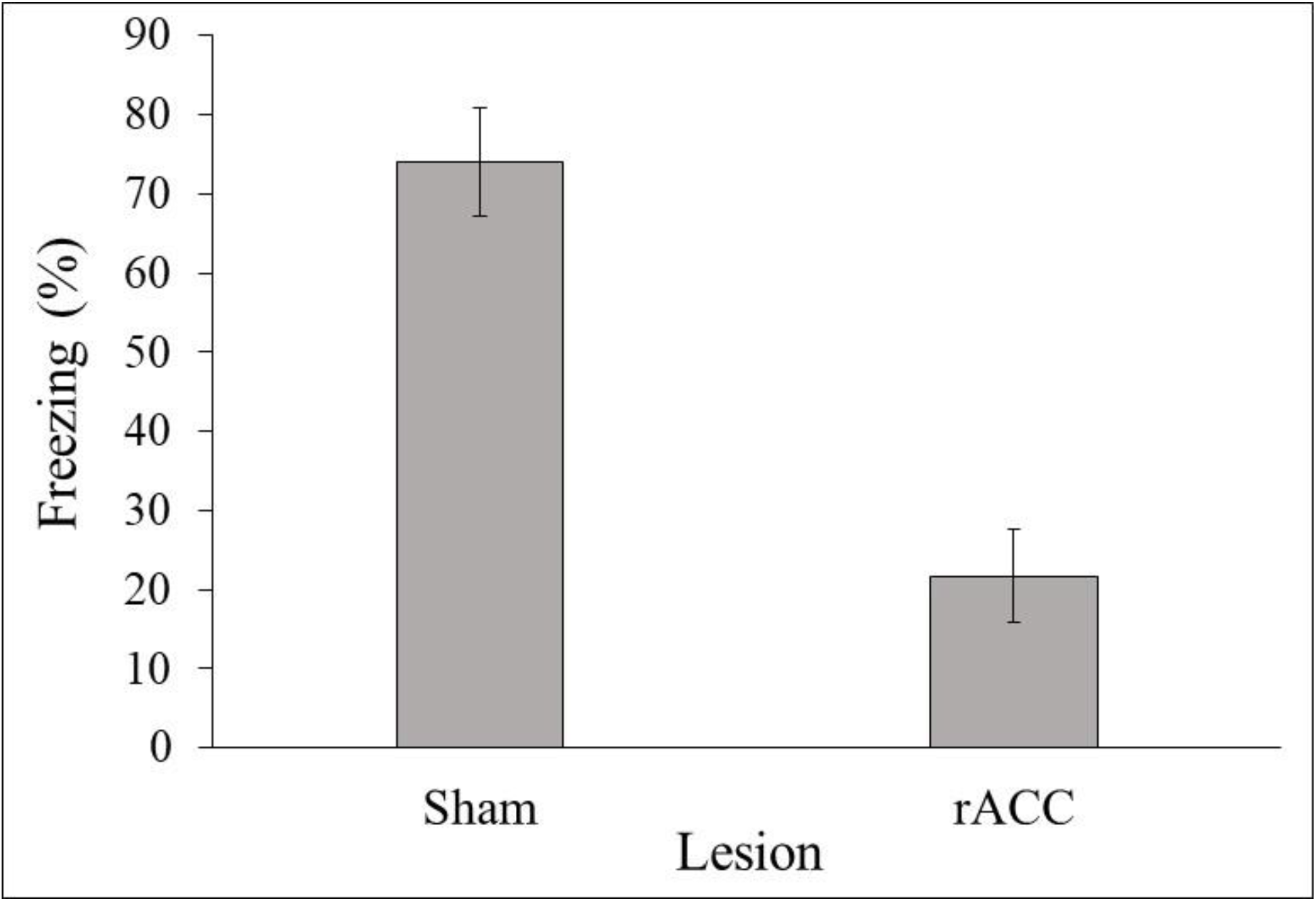
Mean (± SEM) freezing in sham- and rACC-lesioned animals immediately after the cessation of dPAG stimulation applied at the escape threshold.

Fig. 4 presents the mean (± SEM) percentage of time spent freezing in the sham and rACC lesion groups during the contextual fear conditioning test session. Animals with rACC lesions displayed less freezing behavior than sham-lesioned control animals (*t*_18_ = 3.2, *p* < 0.01). Fig. 5 presents the mean (± SEM) percentage of time of formalin-induced behavior in the sham and rACC lesion groups. Animals that received lesions of the rACC displayed a larger amount of formalin-induced behavior than sham-lesioned animals (*t*_18_ = 5.2; *p* < 0.001).

**Figure 4.**
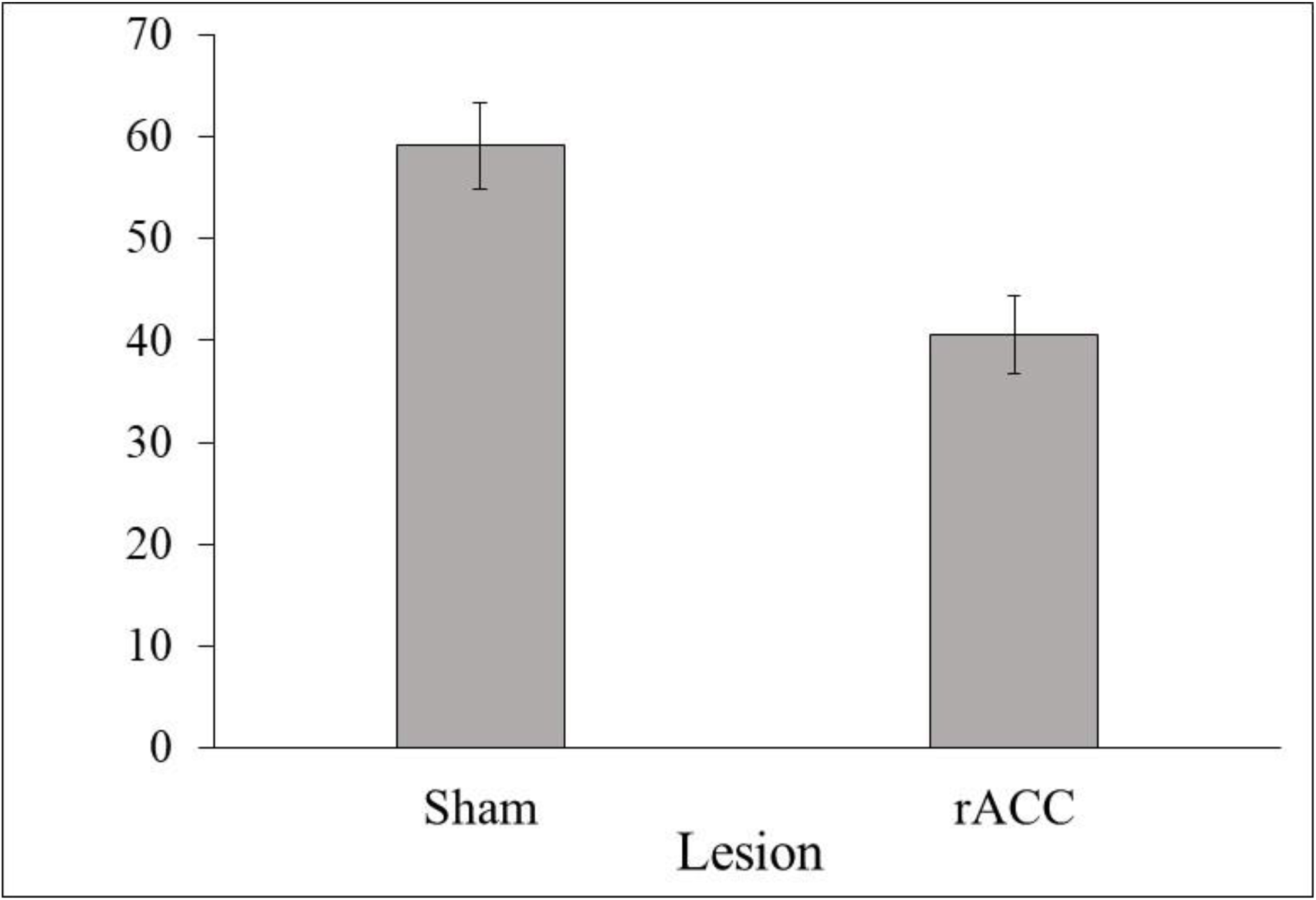
Mean (± SEM) freezing in sham- and rACC-lesioned animals during the contextual fear conditioning test session.

**Figure 5.**
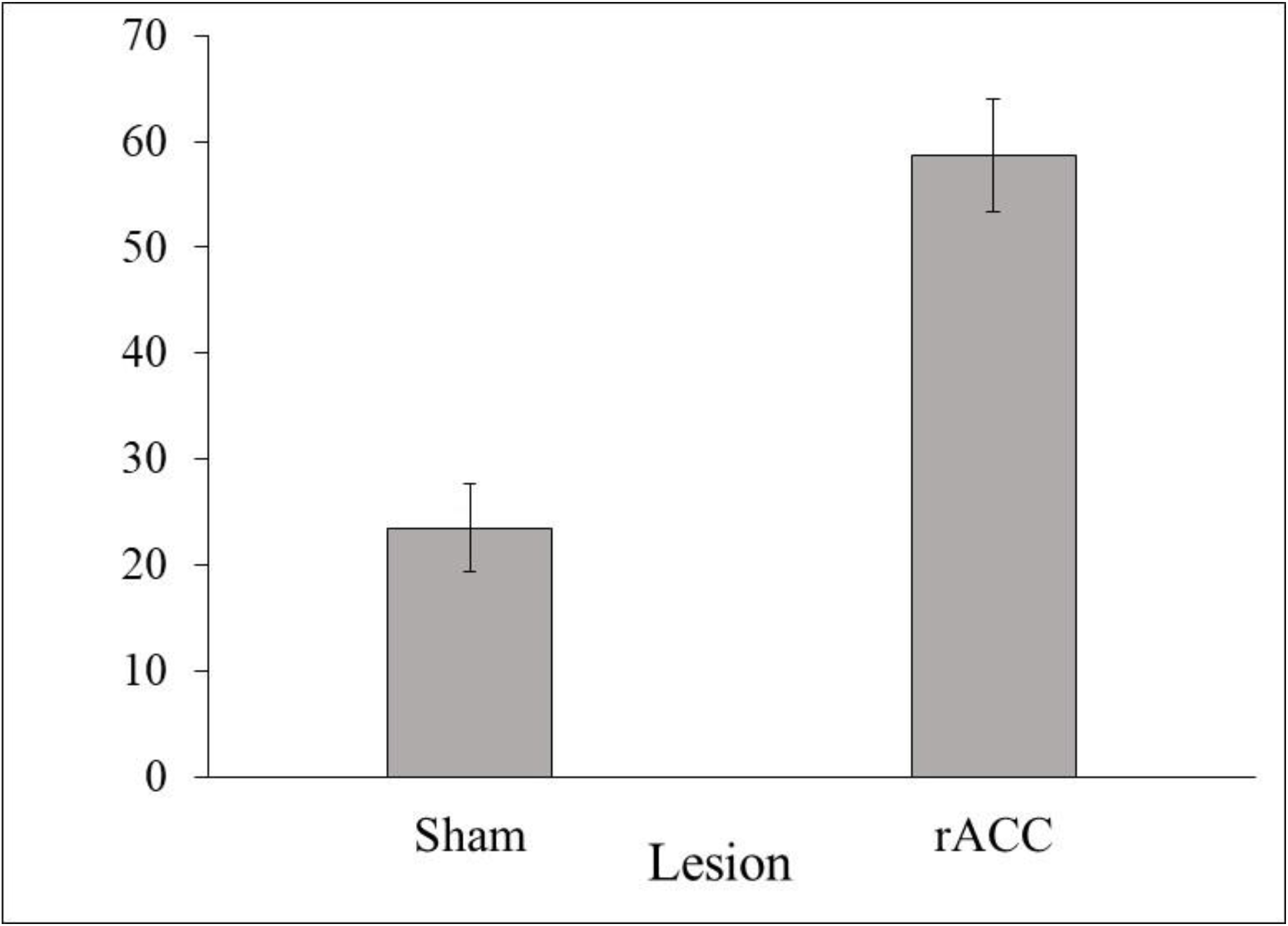
Mean (± SEM) formalin-induced nociceptive behavior in sham- and rACC-lesioned animals during the contextual fear conditioning test session.

## 4. Discussion

Several studies have shown that the dPAG is one of the main substrates of aversion in the brain. Electrical or chemical stimulation of the dPAG produces a set of fear-like responses, such as cardiovascular changes that include an increase in heart rate and blood pressure, hyperventilation, freezing, and active patterns of aversive behavior [19]. The present results indicated that bilateral electrolytic lesions of the rACC did not change the threshold of the electrical current need to elicit freezing or escape responses when applied to the dPAG. These results are consistent with previous reports that showed that defensive freezing and escape responses induced by electrical stimulation of the dPAG did not depend of telencephalic structures, such as the amygdaloid complex [10, 11]. Defensive behavior generated at the level of the dPAG is suggested to be mediated by descending output projections to more caudal brainstem structures that are involved in the motor performance of these defensive responses. Thus, the activation of aversive brain structures closer to motor outputs, such as the dPAG, appears to trigger immediate defensive responses independently from the influence of upstream brain structures.

Importantly, our results also indicated that lesions of the rACC reduced the amount of freezing behavior observed after the interruption of the electrical stimulation of the dPAG at the escape threshold. Therefore, defensive freezing behavior that persisted after the termination of the electrical stimulation of the dPAG encompassed ascending projections that reached the rACC, whereas aversive responses induced by direct dPAG stimulation might exclusively recruit descending projections to motor outputs. Indeed, the dPAG sends projections to the parafascicular nucleus of the intralaminar thalamus [20], which is implicated in the processing of pain. The parafascicular nucleus, in turn, sends projections throughout the anterior cingulate cortex [21, 22].

Studies have shown that the rACC is a crucial region in the regulation of aversive behavior [23, 24]. In the present study, electrolytic lesions of the rACC disrupted post-dPAG stimulation freezing, suggesting that this cortical region is also implicated in the neural circuitry involved in aversive brain stimulation at the level of the dPAG. The present results also showed that rACC lesions markedly reduced the freezing response to contextual cues previously associated with footshock. This finding is consistent with other studies that found that the rACC is involved in the memory process of aversive footshock [12, 13] and contextual fear conditioning [14].

Evidence indicates that freezing in response to contextual cues previously associated with footshock and post-dPAG stimulation freezing might be related to distinct functional systems [19]. Freezing in response to contextual cues depends on a previous association with the unconditioned aversive stimulus (i.e., footshock), whereas post-dPAG stimulation freezing behavior is not context-dependent [25]. The fact that the latter but not the former is insensitive to a contextual shift suggests that post-dPAG stimulation freezing is naturally unconditioned. Electrolytic lesions of the rACC disrupted both types of freezing, indicating that the rACC integrates sensory information to allow the recognition of distinct forms of threatening stimuli.

Our results also demonstrated that rACC lesions substantially increased the amount of formalin-induced behavior in rats exposed to contextual cues previously associated with footshock. Analgesia is inferred from the suppression of nociceptive behavior induced by formalin injection. Thus, the expression of conditioned analgesia was disrupted in rACC-lesioned animals compared with sham-lesioned controls. Consistent with previous results that indicated that the rACC contributes to the inhibitory modulation of nociceptive processing [15], this result provides evidence that the rACC plays an important role in conditioned antinociceptive systems, in addition to contextual and post-dPAG stimulation freezing.

Similar to the effects of lesions of the rACC observed in the present study, electrolytic lesions of the amygdaloid complex also produced a disruptive effect on defensive freezing and conditional analgesia in response to contextual cues previously associated with footshock in the formalin test [26]. Moreover, the blockade of amygdaloid complex activity with muscimol disrupted post-dPAG stimulation freezing did not affect freezing or escape responses triggered by electrical stimulation of the dPAG [10]. Such similar results strongly suggest a close relationship between these two forebrain structures in the modulation of both conditioned analgesia and defensive freezing behavior in response to aversive stimuli. Indeed, the rACC and amygdaloid complex share extensive reciprocal anatomical connections [27–29]. Future studies will clarify how these functional and anatomical relationships between the rACC and amygdaloid complex and their relationship with the dPAG might interact during aversive stimulus processing.

## Acknowledgments

This research was supported by CNPq and FAPERJ grants awarded to JLF. SM was supported by a postdoctoral fellowship from FAPERJ. BOG and AF were supported by graduate student fellowships from CNPq.

